# GenomeChronicler: The Personal Genome Project UK Genomic Report Generator Pipeline

**DOI:** 10.1101/2020.01.06.873026

**Authors:** José Afonso Guerra-Assunção, Lucia Conde, Ismail Moghul, Amy P. Webster, Simone Ecker, Olga Chervova, Christina Chatzipantsiou, Pablo P. Prieto, Stephan Beck, Javier Herrero

**Affiliations:** Infection and Immunity, University College London, London, United Kingdom; Bill Lyons Informatics Centre, UCL Cancer Institute, University College London, London, United Kingdom; Medical Genomics, UCL Cancer Institute, University College London, London, United Kingdom; Lifebit, 9 Appold St, EC2A 2AP London, United Kingdom

**Keywords:** Personal Genomics, PGP-UK, Genomic Report, Open Consent, Participant Engagement, Open Source, Cloud Computing

## Abstract

In recent years, there has been a significant increase in whole genome sequencing data of individual genomes produced by research projects as well as direct to consumer service providers. While many of these sources provide their users with an interpretation of the data, there is a lack of free, open tools for generating reports exploring the data in an easy to understand manner.

GenomeChronicler was developed as part of the Personal Genome Project UK (PGP-UK) to address this need. PGP-UK provides genomic, transcriptomic, epigenomic and self-reported phenotypic data under an open-access model with full ethical approval. As a result, the reports generated by GenomeChronicler are intended for research purposes only and include information relating to potentially beneficial and potentially harmful variants, but without clinical curation.

GenomeChronicler can be used with data from whole genome or whole exome sequencing, producing a genome report containing information on variant statistics, ancestry and known associated phenotypic traits. Example reports are available from the PGP-UK data page (personalgenomes.org.uk/data).

The objective of this method is to leverage existing resources to find known phenotypes associated with the genotypes detected in each sample. The provided trait data is based primarily upon information available in SNPedia, but also collates data from ClinVar, GETevidence and gnomAD to provide additional details on potential health implications, presence of genotype in other PGP participants and population frequency of each genotype.

The analysis can be run in a self-contained environment without requiring internet access, making it a good choice for cases where privacy is essential or desired: any third party project can embed GenomeChronicler within their off-line safe-haven environments. GenomeChronicler can be run for one sample at a time, or in parallel making use of the Nextflow workflow manager.

The source code is available from GitHub (https://github.com/PGP-UK/GenomeChronicler), container recipes are available for Docker and Singularity, as well as a pre-built container from SingularityHub (https://singularity-hub.org/collections/3664) enabling easy deployment in a variety of settings. Users without access to computational resources to run GenomeChronicler can access the software from the Lifebit CloudOS platform (https://lifebit.ai/cloudos) enabling the production of reports and variant calls from raw sequencing data in a scalable fashion.

## 1 Introduction

The publication of the first draft human genome sequence <u>(International Human Genome Sequencing Consortium 2001)</u> promised a revolution in knowledge of how we see ourselves as individuals and how future medical care should take our genetic background into account. Almost ten years later, the perspective of widespread personal genomics was still to be achieved (Venter 2010).

Following the establishment of 23andMe and others from 2007 onwards, there is now a wide range of easily accessible clinical and non-clinical genetic tests that are routinely employed to detect individuals’ carrier status for certain disease genes or particular mutations of clinical relevance.

Many more associations between genotype and phenotype have been highlighted by research, sometimes with uncertain clinical relevance or simply describing personal traits such as eye color (Pontikos et al. 2017; Kuleshov et al. 2019).

Over the past few years, we have seen a dramatic reduction of the cost to sequence the full human genome. This reduction in cost enables many more projects to start using whole genome sequencing (WGS) approaches, as well as the marked rise in the number of personal genomes being sequenced.

Personal genomics is very much a part of the public consciousness as can be seen by the rampant rise in direct to consumer (DTC) genomic analysis offerings on the market. In this context, it is unsurprising that the analysis of one’s own genome provides a valuable educational opportunity (Salari et al. 2013; Linderman et al. 2018) as well as increasing participant engagement as part of biomedical trials (Sanderson et al. 2016).

The Personal Genome Project (PGP) set up by George Church in 2005 is the earliest initiative enabled by the increased popularity of whole genome sequencing and its lowering costs. The global PGP network currently consists of 5 projects spread around the world, managed independently but joined by a common goal of providing open access data containing genomic, environmental and trait information (https://www.personalgenomes.org/).

Data analysis within PGP-UK poses important ethical challenges, as all the data and genome reports are intended to become freely and openly available on the World Wide Web. However, until the completion and approval of the reports, the data must be treated as confidential private information. Prior to enrollment, all participants are well informed through an online study guide and tested for their understanding of the potential risks of participating in a project of this nature. Upon receipt of their report, participants have a cool-off period of four weeks to explore their data and reports and to seek all the required clarifications. During that time, they can trigger the release of their report and data themselves by selecting the ‘release immediately’ option in their personal accounts. To date, 67% of participants have selected this release option. They also have the option to withdraw from the study in which case no release occurs and all data will be deleted. This option has never been selected by any participant. If neither of these options are chosen, the data and reports are released automatically by the end of the cool-off period.

There are several resources aimed at users of DTC genetic testing companies on the internet including Promethease (‘Promethease’ 2019) and Genomelink (‘Genomelink | Upload Raw DNA Data for Free Analysis On 25 Traits’ 2019). There are some other tools with a focus on clinical aspects or particular diseases (Nakken et al. 2018), as well as academic databases containing genotypes of other individuals (Greshake et al. 2014), pharmacogenomic information (Klein and Ritchie 2018) or genotype to phenotype links (Ramos et al. 2014; Pontikos et al. 2017; Kuleshov et al. 2019) that can be useful for the interpretation of personal genomes. Many of these are linked into resources like SNPedia (Cariaso and Lennon 2012), allowing a wide range of exploration options for the known associations of each genotype from multiple perspectives.

Surprisingly, we found no pre-existing solution that would allow the annotation and evaluation of variants on the whole genome level, assessment of ancestry and more focused analysis of variants that have been previously associated with specific phenotypes. In particular, one that could be run locally ensuring full control of the data before the results are scrutinized and approved.

GenomeChronicler represents, to the best of our knowledge, the first pipeline that can be run offline or in the cloud, to generate personal genomics reports that are not limited to disease only, from whole genome or whole exome sequencing data.

GenomeChronicler contains a database of positions of interest for ancestry or phenotype. The genotype at each of these positions is inferred from the user provided data that has been mapped to the human genome. These genotypes are then compared to local versions of a series of publicly available resources to infer ancestry and likely phenotypes for each individual participant. These results are then presented as a PDF document containing hyperlinks where more information about each variant and phenotype can be found. A visual representation of the pipeline and its underlying resources is shown in Figure 1.

**Figure 1:**
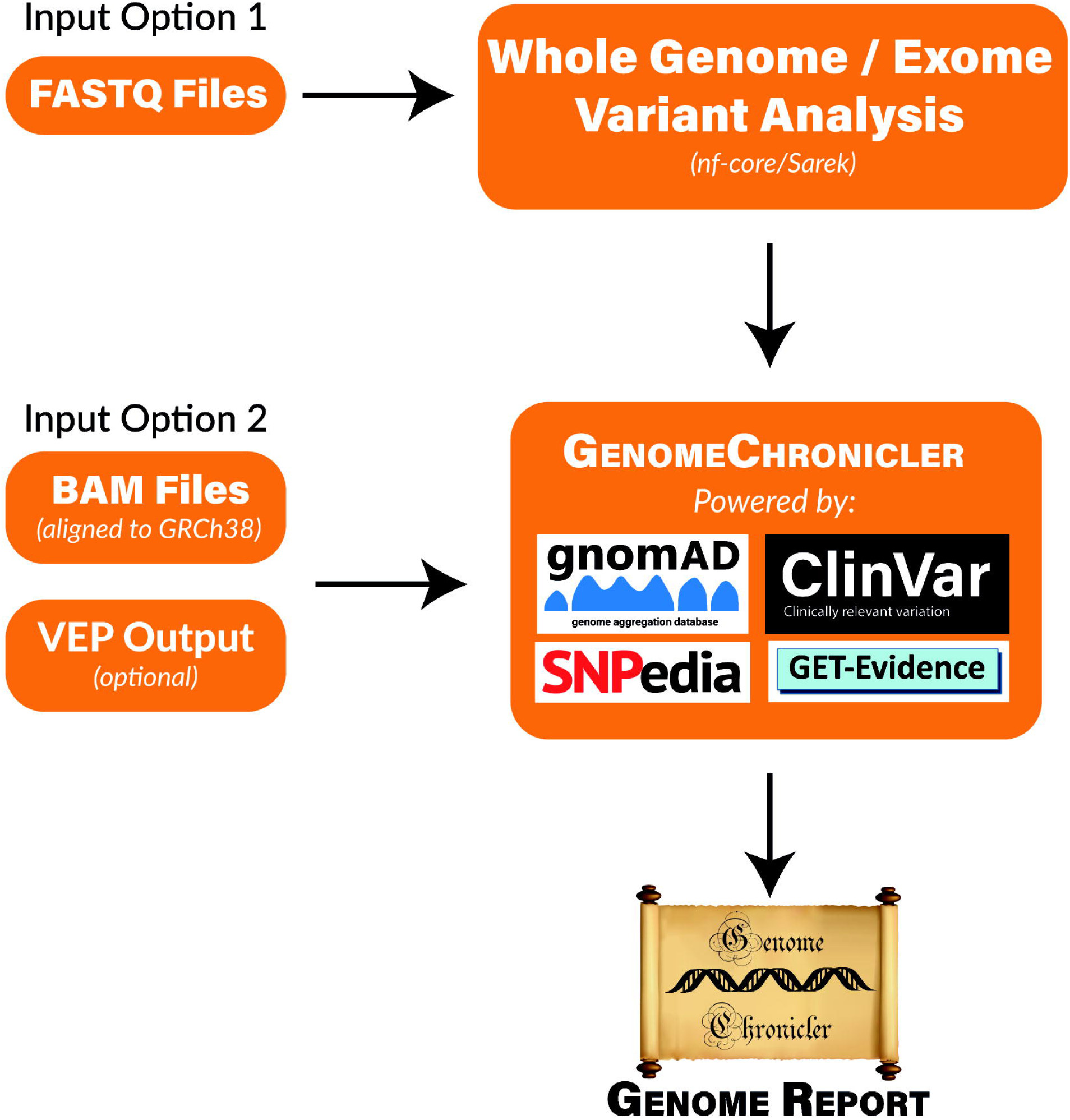
Flow Diagram of GenomeChronicler processing pipeline, illustrating the multiple entry points for the pipeline, resources integrated by default and generated outcomes. Either entry point of the pipeline can be run locally in a single machine, as a Nextflow workflow or in the Cloud. All source code and integrations are freely available in their respective GitHub repositories. The stand-alone GenomeChronicler is available at (https://github.com/PGP-UK/GenomeChronicler), the integration of GenomeChronicler with Nextflow is available at (https://github.com/PGP-UK/GenomeChronicler-nf) and the combined GenomeChronicler with Sarek variant calling is available at (https://github.com/PGP-UK/GenomeChronicler-Sarek-nf). The recipe files for the Docker and Singularity containers are available within the respective GitHub repositories. The resource logos are reproduced from the respective resource websites and remain copyright of their original owner.

This pipeline will continue to be improved and expanded by PGP-UK, e.g. to include methylome and transcriptome reports (Beck et al. 2018). We envision this project will also be useful to other research endeavors that want to provide personal genomes information to their participants to increase engagement; e.g. to altruistic individuals who have obtained their whole genome sequencing data from a DTC or health care provider and are looking for an ethics-approved framework to share their data. PGP-UK already supports this through their Genome Donation programme.

## 2 Materials and Methods

### 2.1 Data Input

The GenomeChronicler pipeline was designed to run downstream of a standardised germline variant calling pipeline. GenomeChronicler requires a pre-processed BAM or CRAM file with deduplicated and quality recalibrated alignments against the GRCh38 genome assembly and optionally, the summary HTML report produced by the Ensembl Variant Effect Predictor (McLaren et al. 2016).

GenomeChronicler can be run with any variant caller provided that the reference dataset is matched to the reference genome used (the included GenomeChronicler databases currently use GRCh38). It is also imperative, to obtain good quality results, that the BAM or CRAM files used have had their duplicates removed and quality recalibrated prior to being used for GenomeChronicler.

To simplify this entire process and to make the tool more accessible to users who may not know how to run a germline variant calling pipeline, GenomeChronicler can also be run in a fully automated mode from the raw sequencing data, where the germline variant calling pipeline is also run and the whole process is managed by the Nextflow workflow management system <u>(Di Tommaso et al. 2017)</u>. In this scenario, GenomeChronicler uses the Sarek pipeline (https://github.com/nf-core/sarek, Garcia et al. 2020) to process raw FASTQ files in a manner that follows the GATK variant calling best practices guidelines (Van der Auwera et al. 2013). Manual inspection of the initial quality control steps of Sarek is recommended prior to perusing the final results.

The combined version of Sarek + GenomeChronicler written using the Nextflow workflow manager (Di Tommaso et al. 2017) is available both on Github (https://github.com/PGP-UK/GenomeChronicler-Sarek-nf) and on Lifebit CloudOS.

### 2.2 Ancestry Inference

We infer the ancestry of each individual through a Principal Components Analysis (PCA) which is a widely used approach for identifying ancestry similarities among individuals (Novembre et al. 2008). For each sample of interest, we intersect the genotypes with a reference dataset consisting of genotypes from the 1000 Genomes Project samples (The 1000 Genomes Project Consortium 2015), containing individuals from 26 different worldwide populations and applying PCA on the resulting genotype matrix.

The reference samples from the 1000 Genome Project are filtered to keep only unrelated individuals. In order to avoid strand issues when merging the datasets, all ambiguous (A/T and C/G) SNPs were removed, as well as non-biallelic SNPs, SNPs with >5% of missing data, rare variants (MAF < 0.05) and SNPs out of Hardy-Weinberg equilibrium (pval < 0.0001). From the remaining SNPs, a subset of unlinked SNPs are selected by pruning those with r2 > 0.1 using 100-SNP windows shifted at 5-SNP intervals.

These genotypes are used to run PCA based on the variance-standardized relationship matrix, selecting twenty as the number of PCs to be extracted. We then project the data over the first three principal components to identify clusters of populations and highlight the sample of unknown ancestry on the resulting plot.

Here, we used PLINK (Purcell et al. 2007) to process the genotype data and the R Statistical Computing platform for plotting the final PCA figures to illustrate the ancestry of each sample. An example of the distribution of the reference samples on the PCA is shown in Figure 2.

**Figure 2:**
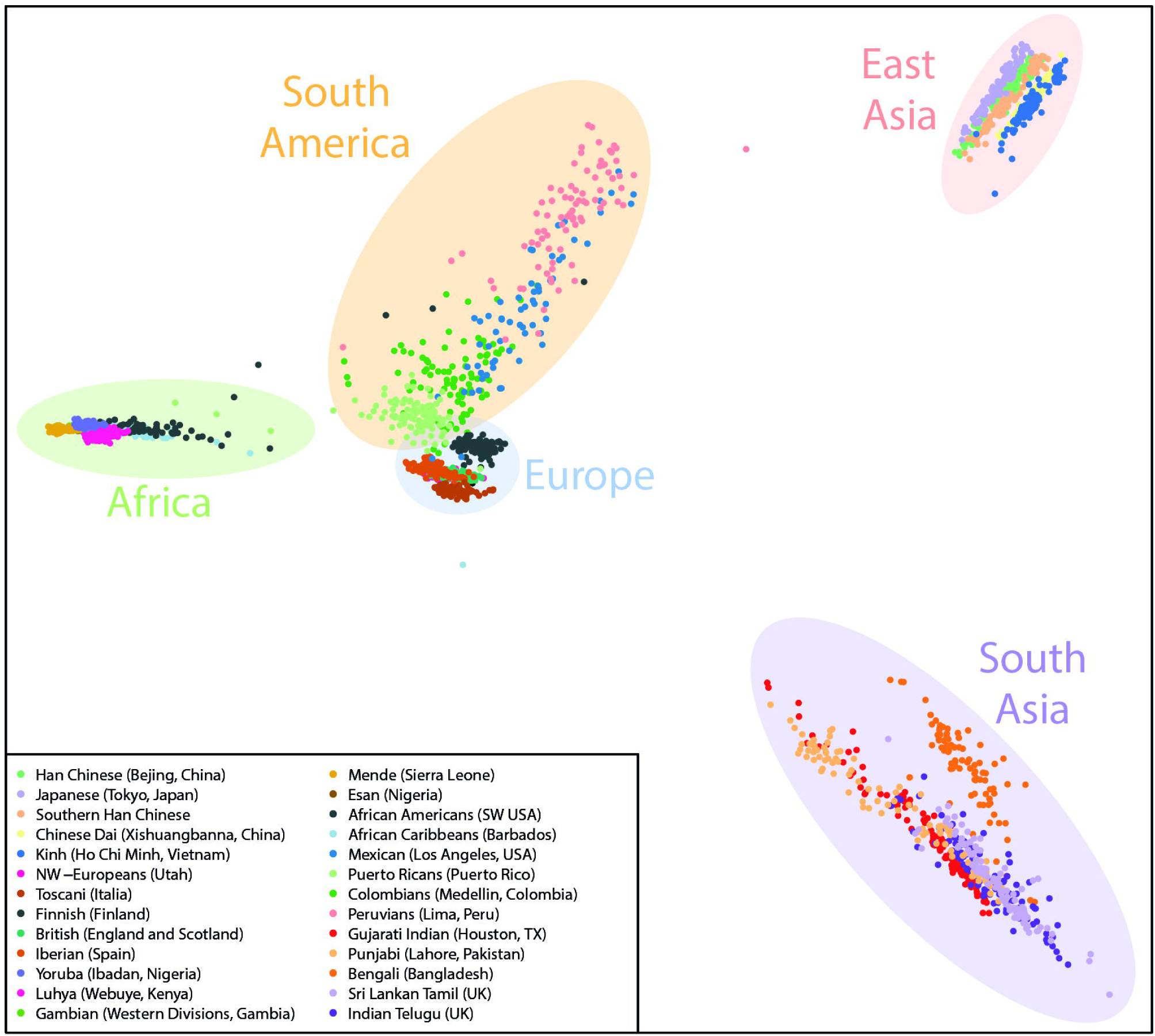
Example Ancestry PCA plot containing the current reference data from the 1000 genomes project used by GenomeChronicler, with shaded areas broadly illustrating the origin of the populations represented.

### 2.3 Variant Annotation Databases

#### 2.3.1 SNPedia

SNPedia (Cariaso and Lennon 2012) is a large public repository of manually added as well as automatically mined genotype to phenotype links sourced from existing literature. SNPedia is the core resource behind the phenotype tables in GenomeChronicler; it provides annotations for both single-gene phenotypes as well as for a few phenotypes involving multiple loci referred to as genosets in the produced reports.

#### 2.3.2 ClinVar

ClinVar (Landrum and Kattman 2018) is a database hosted by the NCBI that focuses exclusively on variants related to health and has been running since 2013. In comparison to SNPedia, ClinVar is a much smaller database but it is closely linked to the clinical relevance of each variant. ClinVar is curated more strictly with a clinical review – something unique among the data sources used by GenomeChronicler.

#### 2.3.3 GETevidence

GETevidence was developed as part of the Personal Genome Project Harvard (Mao et al. 2016) to showcase the variants present within its participants and to allow manual annotation and interpretations of the results. For some of the genotypes present, it also contains manual annotations that have been added by the users or curation team. GETevidence allows individuals to compare their genotypes with those from other personal genomes available within the PGP-Harvard project.

#### 2.3.4 gnomAD

Spanning several human populations, the Genome Aggregation Database (gnomAD) (Karczewski et al. 2019) aggregates data from multiple sources to produce an atlas of variation across the human genome. Extensively annotated and now covering most of the latest assembly of the human genome, these links enable easy access to information such as allele frequencies for the genotype across different populations around the world, as well as some annotation context for each variant, regarding potential effect on genes if relevant and how selection forces are constraining the genomic locus.

### 2.4 Database Availability, Building and Update

The underlying databases required to run GenomeChronicler are provided within the package. A set of scripts to regenerate these SQLite databases is also provided within the source code. The datasets are limited to positions of interest is compiled so that when genotyping is performed only relevant positions are computed to save computational time.

SNPedia provides an API to query its records in a systematic way. The other linked databases provide regular dumps of the whole dataset, enabling easy assessment for which dbSNP rs identifiers are represented within the full database. The use of rs identifiers and genotypes to link between the different databases enables an unambiguous way to compare information between different resources.

### 2.5 Genotype assessment and reporting

Typical germline variant calling pipelines result in a VCF file where positions that match the reference sequence are not reported. Homozygous reference genotypes thus become indistinguishable from positions in the genome where there is no read coverage.

To ensure comparable results between runs, genotype VCFs (gVCFs) instead of VCFs are computed during each run of GenomeChronicler, but only for a subset of genomic positions that informative for ancestry inference or phenotype annotation, saving computational time and storage space.

### 2.6 The Genome Report Template

GenomeChronicler is designed in a modular way where the final report is only compiled at the end, integrating all the results. To customize the report layout, the content and the amount of extra information, GenomeChronicler uses a template file written in LaTeX. For example, one can modify the branding and introductory text of the report, integrate custom or third-party analyses provided the results are in a format that can be typeset using LaTeX, omit certain sections, or even modify the structure of the report produced.

### 2.7 Output Files

The main output of GenomeChronicler is a report in PDF format, containing information from all sections of the pipeline that have run as set by the LaTeX template provided when running the script. Additionally, an Excel file containing the genotype phenotype link information, and all corresponding hyperlinks is also produced, allowing the user to explore the results in a familiar environment. While most intermediate files are automatically removed at the end of the GenomeChronicler run, the original PDF version of the ancestry PCA plot, as well as a file containing the sample name, genotyping results and pipeline log files are retained within the results directory to ease automation.

### 2.8 Pipeline Validation

To further validate the pipeline, 1000 Genome Project generated illumina data for sample NA12878 was used. Genomic data for sample NA12878 mapped to the human reference genome (GRCh38) was retrieved from the 1000 Genome Project (ftp://ftp.1000genomes.ebi.ac.uk/vol1/ftp/data_collections/1000_genomes_project/data/CEU/NA12878/alignment/) and converted to BAM file using the SAMtools toolkit. High confidence genotype calls were retrieved from Genome-in-a-Bottle (ftp://ftp-trace.ncbi.nlm.nih.gov/giab/ftp/release/NA12878_HG001/latest/GRCh38/). The GenomeChronicler pipeline was run on the data, and the resulting genotype calls in high confidence regions were compared to the reference calls using BCFtools to assess genotype concordance.

### 2.9 Running GenomeChronicler

In line with the PGP-UK data, all the code for GenomeChronicler is freely available. To make it easier to implement, several options are available to eliminate the need for installing dependencies and underlying packages, or even the need to have access to computer hardware capable of handling the processing of a human genome. The range of options available is detailed below and illustrated in Figure 1.

#### 2.9.1 Running GenomeChronicler Locally

##### 2.9.1.1 From the available source code

The source code for GenomeChronicler is available on GitHub at https://github.com/PGP-UK/GenomeChronicler. A setup script is included to automatically download the pre-compiled accessory databases and other required data. Software dependencies including LaTeX, R and Perl need to be installed independently if not using the Singularity container. The provided Singularity recipe file provides a useful list of required packages, in particular for those installing it on a Debian/Ubuntu based system.

##### 2.9.1.2 Using a pre-compiled container

GenomeChronicler is also available as a Singularity container (Kurtzer, Sochat, and Bauer 2017) with all dependencies pre-installed and ready. This can be obtained from SingularityHub (Sochat, Prybol, and Kurtzer 2017) by running the command: `singularity pull shub://PGP-UK/GenomeChronicler` on any machine that has Singularity installed.

Once downloaded, the main script (GenomeChronicler_mainDruid.pl) can be run with the desired data and options to produce genome reports.

#### 2.9.2 Running GenomeChronicler on Cloud

To enable reproducible, massively parallel, cloud native analyses, GenomeChronicler has also been implemented as a Nextflow pipeline. The implementation abstracts the installation overhead from the end user, as all the dependencies are already available via pre-built containers, integrated seamlessly in the Nextflow pipeline. The source code for this implementation is available on GitHub at https://github.com/PGP-UK/GenomeChronicler-nf, as a standalone Nextflow process.

To provide an end-to-end FASTQ to PGP-UK reports pipeline, we also implemented an integration of GenomeChronicler, with a curated and widely used by the bioinformatics community pipeline, namely Sarek (Garcia et al. 2020; Ewels et al. 2019). This PGP-UK implementation of Sarek is available on GitHub at https://github.com/PGP-UK/GenomeChronicler-Sarek-nf. The aforementioned pipeline is available in the collection of curated pipelines on the Lifebit CloudOS platform (https://cloudos.lifebit.ai/). Lifebit CloudOS enables users without any prior cloud computing knowledge to deploy analysis in the cloud. In order to run the pipeline, the user only needs to specify input files, desired parameters and select resources from an intuitive graphical user interface. After the completion of the analysis on Lifebit CloudOS, the user has a permanent shareable live link that includes performance and file metadata, the associated GitHub repository revision and also links to the generated results. The relevant analysis page can be used to repeat the exact same analysis. The analysis page for the PGP-UK user with id uk35C650 can be accessed in the following permalink https://cloudos.lifebit.ai/public/jobs/5e3582dae3474100f4665c7a. Each analysis can have different privacy settings allowing the user to choose if the results are publicly visible, making it easier for sharing or private use, thus maintaining data confidentiality.

## 3 Results

The main resulting document is a PDF file which contains sections related to variants of unknown significance, ancestry estimation (as exemplified in Figure 2) and variants with associated phenotypes, separated by either potentially beneficial or potentially harmful phenotypes as well as phenotypes affected by multiple variants, referred to as genosets (Cariaso and Lennon 2012).

Initial versions of the GenomeChronicler pipeline were validated by comparing its results to those provided by DTC company 23andMe for participant PGP-UK1, as well as phenotype feedback from the pilot participants (Beck et al. 2018).

Further validations was done using sample NA12878, which is an often-analysed as a benchmark reference for personal genomics.

The GATK genotype calls produced as part of GenomeChronicler were directly compared to the high confidence variant calling for the sample as part of the Genome-in-a-Bottle consortium (Zook et al. 2014). The concordance rate was 99.97% at the genotype level, resulting in no phenotype changes.

Sample NA12878 is part of pedigree 1463 from the HapMap project and is known to correspond to a female individual of CEPH ancestry. These are correctly reflected in the ancestry and genoset sections of the GenomeChronicler report.

To date, more than one hundred such reports have been produced and made available as part of the PGP-UK (Beck et al. 2018). They are publicly available in the PGP-UK open access data page (https://www.personalgenomes.org.uk/data/). This collection contributes to the educational potential of the project as a whole. On one hand, it allows participants of PGP-UK and other users of the GenomeChronicler tool to compare their genome report results to those of other individuals. On the other hand, it allows individuals that are interested in the subject but did not have their genome sequenced to explore the kind of information that one can learn from a personal genome.

While the method presented here focuses on the analysis of the genomic data (whole genome and whole exome), PGP-UK also contains multi-omics data, including RNAseq and methylation data, as well as genotype data sourced elsewhere (e.g. 23andMe) and deposited by the participants.

Methods such as GenomeChronicler allow other research projects in possession of personal genome data to easily produce genome reports, customise them with static text providing information about the project that can differ from the default template file, or even add links to other relevant databases.

## 4 Conclusions

Here we present GenomeChronicler, a computational pipeline to produce genome reports including variant calling summary data, ancestry inference, and phenotype annotation from genotype data for personal genomics data obtained through whole genome or whole exome sequencing.

The pipeline is modular, fully open source, and available as containers and on the Lifebit CloudOS computing platform, enabling easy integration with other projects, regardless of available computational resources and bioinformatics expertise.

The pipeline presented here incorporates a range of well-established open source resources, which have been validated independently in different scenarios (Garcia et al. 2020). We have also cross-referenced the data produced by this pipeline to ensure it is providing a coherent output (Chervova et al. 2019).

While we follow the GATK best practices, as implemented in Sarek, to produce an accurate and reliable variant call set, unforeseen sources of error can be introduced at the sequencing stage, resulting in the pipeline potentially calling an erroneous genotype at a certain genomic position.

Finally, the interpretation of genotype to phenotype links is heavily context-dependent and fraught with its own challenges. Recognising that this task requires experience and/or cognitive abilities that cannot be imparted on an automated computer system, we instead opted for providing a report that focuses on the biomedical and phenotypic associations obtained through SNPedia (Cariaso and Lennon 2012), supplemented with hyperlinks to a wide range of other databases. This allows the user to explore the results and the supporting research data in more depth if desired. Some of the reported links between genotypes and phenotypes have been strongly validated by multiple research groups over the years, while others are not as well supported, and as such, require careful interpretation by the user.

This work was developed as part of PGP-UK and incorporates feedback from early participants to improve the usefulness of the reports produced, and of participant engagement. It is designed to be easily expandable, adaptable to other contexts and most of all, suitable for projects with a wide range of ethical requirements, from those that need the data to be processed inside a safe-haven environment to those that process all the data in the public domain. It can also be of interest to educational groups such as Open Humans (Greshake Tzovaras et al. 2019). Open Humans (https://www.openhumans.org/) is a vibrant community of researchers, patients, data and citizen scientists who want to learn more about themselves.

For PGP-UK participants, there is a well-established ethical framework that ensures that participants are aware of the limitations of the information they receive. It also makes provision for the project to refrain from issuing reports if the quality of the input data fails the quality control stage.

Personal genomics has become a public commodity and individuals can access their own or even someone else’s genome. It is important to note that GenomeChronicler is essentially a tool that collates information from different sources but is not suitable for the clinical interpretation of the results. Indeed, inaccurate interpretation might result from poor quality genomic data or unreliable annotations. However, the potential for negative consequences should be minimal provided the users heed the stated recommendations of not relying on this tool for clinical decision making.

Future directions for this work will include the integration of other omics data types that are produced within PGP-UK, as well as potentially expanding the databases that are linked by default when running the pipeline.

We hope that GenomeChronicler will be useful to other projects and interested individuals. As it is open source, the pipeline can easily adapt custom templates to satisfy any curiosity-driven analyses and increase the level of genomic understanding in general.

## 5 Conflict of Interest

Pablo P. Prieto is CTO of Lifebit and Christina Chatzipantsiou is an employee of Lifebit. All other authors declare that the research was conducted in the absence of any commercial or financial relationships that could be construed as a potential conflict of interest.

## 6 Author Contributions

J.A.G.-A. led the development and implementation of the method and wrote the manuscript with input from all authors. J.A.G.-A., L.C. contributed computer code. C.C. contributed the Nextflow and Lifebit CloudOS integrations with support from P.P.P.. J.A.G.-A., L.C., I.M., A.P.W., S.E., JH., O.C. and S.B. contributed to the conceptual development of the method and usability. All authors read and approved the manuscript.

## 7 Funding

PGP-UK gratefully acknowledges support from the Frances and Augustus Newman Foundation, Dangoor Education and the National Institute for Health Research (NIHR) UCLH Biomedical Research Centre (BRC369/CN/SB/101310). The views expressed are those of the authors and not necessarily those of the NIHR or the Department of Health and Social Care.

## 8 Acknowledgements

The authors acknowledge the use of the UCL Legion High Performance Computing Facility (Legion@UCL) and associated support services. The authors thank all PGP-UK participants for their contributions to the project and feedback on the items that feature in the reports produced by GenomeChronicler. This manuscript has also been submitted as a preprint to the bioRxiv preprint server, available at https://www.biorxiv.org/content/10.1101/2020.01.06.873026.

## 9 Data Availability Statement

The datasets analyzed and used for the development of the approach here described are deposited at the European Nucleotide Archive (ENA) hosted by the EMBL-EBI under the umbrella accession PRJEB24961. [https://www.ebi.ac.uk/ena/data/view/PRJEB24961]. The PGP-UK pilot data was described in a data descriptor published in Scientific Data (Chervova et al. 2019). The source code for the software is deposited and maintained in GitHub and available at [https://github.com/PGP-UK/GenomeChronicler]. The Nextflow integrated version is available at [https://github.com/PGP-UK/GenomeChronicler-nf] and finally, the version also containing the Sarek variant calling pipeline is available at [https://github.com/PGP-UK/GenomeChronicler-Sarek-nf]. Reports generated using this approach for PGP-UK samples are archived in the PGP-UK data page https://www.personalgenomes.org.uk/data.

## Notes

http://www.personalgenomes.org.uk/data/

### Summary of Updates

Providing updated version after peer review.

## References

Beck, Stephan, Alison M. Berner, Graham Bignell, Maggie Bond, Martin J. Callanan, Olga Chervova, Lucia Conde, et al. 2018. ‘Personal Genome Project UK (PGP-UK): A Research and Citizen Science Hybrid Project in Support of Personalized Medicine’. BMC Medical Genomics 11 (1): 108. https://doi.org/10.1186/s12920-018-0423-1.

Cariaso, Michael, and Greg Lennon. 2012. ‘SNPedia: A Wiki Supporting Personal Genome Annotation, Interpretation and Analysis’. Nucleic Acids Research 40 (Database issue): D1308–1312. https://doi.org/10.1093/nar/gkr798.

Chervova, Olga, Lucia Conde, José Afonso Guerra-Assunção, Ismail Moghul, Amy P. Webster, Alison Berner, Elizabeth Larose Cadieux, et al. 2019. ‘The Personal Genome Project-UK, an Open Access Resource of Human Multi-Omics Data’. Scientific Data 6 (1): 1–10. https://doi.org/10.1038/s41597-019-0205-4.

Di Tommaso, Paolo, Maria Chatzou, Evan W. Floden, Pablo Prieto Barja, Emilio Palumbo, and Cedric Notredame. 2017. ‘Nextflow Enables Reproducible Computational Workflows’. Nature Biotechnology 35 (4): 316–19. https://doi.org/10.1038/nbt.3820.

Ewels, Philip A., Alexander Peltzer, Sven Fillinger, Johannes Alneberg, Harshil Patel, Andreas Wilm, Maxime Ulysse Garcia, Paolo Di Tommaso, and Sven Nahnsen. 2019. ‘Nf-Core: Community Curated Bioinformatics Pipelines’. BioRxiv, May, 610741. https://doi.org/10.1101/610741.

Garcia, Maxime, Szilveszter Juhos, Malin Larsson, Pall I. Olason, Marcel Martin, Jesper Eisfeldt, Sebastian DiLorenzo, et al. 2020. “Sarek: A Portable Workflow for Whole-Genome Sequencing Analysis of Germline and Somatic Variants.” F1000Research 9 (January): 63. https://doi.org/10.12688/f1000research.16665.1.

‘Genomelink | Upload Raw DNA Data for Free Analysis On 25 Traits’. 2019. 2019. https://genomelink.io/.

Greshake, Bastian, Philipp E. Bayer, Helge Rausch, and Julia Reda. 2014. ‘OpenSNP–A Crowdsourced Web Resource for Personal Genomics’. PLOS ONE 9 (3): e89204. https://doi.org/10.1371/journal.pone.0089204.

Greshake Tzovaras, Bastian, Misha Angrist, Kevin Arvai, Mairi Dulaney, Vero Estrada-Galiñanes, Beau Gunderson, Tim Head, et al. 2019. ‘Open Humans: A Platform for Participant-Centered Research and Personal Data Exploration’. GigaScience 8 (6). https://doi.org/10.1093/gigascience/giz076.

Karczewski, Konrad J., Laurent C. Francioli, Grace Tiao, Beryl B. Cummings, Jessica Alföldi, Qingbo Wang, Ryan L. Collins, et al. 2019. ‘Variation across 141,456 Human Exomes and Genomes Reveals the Spectrum of Loss-of-Function Intolerance across Human Protein-Coding Genes’. BioRxiv, August, 531210. https://doi.org/10.1101/531210.

Klein, Teri E., and Marylyn D. Ritchie. 2018. ‘PharmCAT: A Pharmacogenomics Clinical Annotation Tool’. Clinical Pharmacology and Therapeutics 104 (1): 19–22. https://doi.org/10.1002/cpt.928.

Kuleshov, Volodymyr, Jialin Ding, Christopher Vo, Braden Hancock, Alexander Ratner, Yang Li, Christopher Ré, Serafim Batzoglou, and Michael Snyder. 2019. ‘A Machine-Compiled Database of Genome-Wide Association Studies’. Nature Communications 10 (1): 1–8. https://doi.org/10.1038/s41467-019-11026-x.

Kurtzer, Gregory M., Vanessa Sochat, and Michael W. Bauer. 2017. ‘Singularity: Scientific Containers for Mobility of Compute’. PLOS ONE 12 (5): e0177459. https://doi.org/10.1371/journal.pone.0177459.

Landrum, Melissa J., and Brandi L. Kattman. 2018. ‘ClinVar at Five Years: Delivering on the Promise’. Human Mutation 39 (11): 1623–30. https://doi.org/10.1002/humu.23641.

Linderman, Michael D., Saskia C. Sanderson, Ali Bashir, George A. Diaz, Andrew Kasarskis, Randi Zinberg, Milind Mahajan, Sabrina A. Suckiel, Micol Zweig, and Eric E. Schadt. 2018. ‘Impacts of Incorporating Personal Genome Sequencing into Graduate Genomics Education: A Longitudinal Study over Three Course Years’. BMC Medical Genomics 11 (1): 5. https://doi.org/10.1186/s12920-018-0319-0.

Mao, Qing, Serban Ciotlos, Rebecca Yu Zhang, Madeleine P. Ball, Robert Chin, Paolo Carnevali, Nina Barua, et al. 2016. ‘The Whole Genome Sequences and Experimentally Phased Haplotypes of over 100 Personal Genomes’. GigaScience 5 (1): 42. https://doi.org/10.1186/s13742-016-0148-z.

McLaren, William, Laurent Gil, Sarah E. Hunt, Harpreet Singh Riat, Graham R. S. Ritchie, Anja Thormann, Paul Flicek, and Fiona Cunningham. 2016. ‘The Ensembl Variant Effect Predictor’. Genome Biology 17 (1): 122. https://doi.org/10.1186/s13059-016-0974-4.

Nakken, Sigve, Ghislain Fournous, Daniel Vodák, Lars Birger Aasheim, Ola Myklebost, and Eivind Hovig. 2018. ‘Personal Cancer Genome Reporter: Variant Interpretation Report for Precision Oncology’. Bioinformatics (Oxford, England) 34 (10): 1778–80. https://doi.org/10.1093/bioinformatics/btx817.

Novembre, John, Toby Johnson, Katarzyna Bryc, Zoltán Kutalik, Adam R. Boyko, Adam Auton, Amit Indap, et al. 2008. ‘Genes Mirror Geography within Europe’. Nature 456 (7218): 98–101. https://doi.org/10.1038/nature07331.

Pontikos, Nikolas, Jing Yu, Ismail Moghul, Lucy Withington, Fiona Blanco-Kelly, Tom Vulliamy, Tsz Lun Ernest Wong, et al. 2017. ‘Phenopolis: An Open Platform for Harmonization and Analysis of Genetic and Phenotypic Data’. Bioinformatics 33 (15): 2421–23. https://doi.org/10.1093/bioinformatics/btx147.

‘Promethease’. 2019. 2019. https://www.promethease.com/.

Purcell, Shaun, Benjamin Neale, Kathe Todd-Brown, Lori Thomas, Manuel A. R. Ferreira, David Bender, Julian Maller, et al. 2007. ‘PLINK: A Tool Set for Whole-Genome Association and Population-Based Linkage Analyses’. American Journal of Human Genetics 81 (3): 559–75. https://doi.org/10.1086/519795.

Ramos, Erin M, Douglas Hoffman, Heather A Junkins, Donna Maglott, Lon Phan, Stephen T Sherry, Mike Feolo, and Lucia A Hindorff. 2014. ‘Phenotype–Genotype Integrator (PheGenI): Synthesizing Genome-Wide Association Study (GWAS) Data with Existing Genomic Resources’. European Journal of Human Genetics 22 (1): 144–47. https://doi.org/10.1038/ejhg.2013.96.

Salari, Keyan, Konrad J. Karczewski, Louanne Hudgins, and Kelly E. Ormond. 2013. ‘Evidence That Personal Genome Testing Enhances Student Learning in a Course on Genomics and Personalized Medicine’. PloS One 8 (7): e68853. https://doi.org/10.1371/journal.pone.0068853.

Sanderson, Saskia C., Michael D. Linderman, Sabrina A. Suckiel, George A. Diaz, Randi E. Zinberg, Kadija Ferryman, Melissa Wasserstein, Andrew Kasarskis, and Eric E. Schadt. 2016. ‘Motivations, Concerns and Preferences of Personal Genome Sequencing Research Participants: Baseline Findings from the HealthSeq Project’. European Journal of Human Genetics 24 (1): 14–20. https://doi.org/10.1038/ejhg.2015.118.

Sochat, Vanessa V., Cameron J. Prybol, and Gregory M. Kurtzer. 2017. ‘Enhancing Reproducibility in Scientific Computing: Metrics and Registry for Singularity Containers’. PLOS ONE 12 (11): e0188511. https://doi.org/10.1371/journal.pone.0188511.

The 1000 Genomes Project Consortium. 2015. ‘A Global Reference for Human Genetic Variation’. Nature 526 (7571): 68–74. https://doi.org/10.1038/nature15393.

Van der Auwera, Geraldine A., Mauricio O. Carneiro, Chris Hartl, Ryan Poplin, Guillermo del Angel, Ami Levy-Moonshine, Tadeusz Jordan, et al. 2013. ‘From FastQ Data to High Confidence Variant Calls: The Genome Analysis Toolkit Best Practices Pipeline’. Current Protocols in Bioinformatics / Editoral Board, Andreas D. Baxevanis … [et Al.] 11 (1110): 11.10.1–11.10.33. https://doi.org/10.1002/0471250953.bi1110s43.

Venter, J. Craig. 2010. ‘Multiple Personal Genomes Await’. Nature 464 (7289): 676–77. https://doi.org/10.1038/464676a.

Zook, Justin M., Brad Chapman, Jason Wang, David Mittelman, Oliver Hofmann, Winston Hide, and Marc Salit. 2014. “Integrating Human Sequence Data Sets Provides a Resource of Benchmark SNP and Indel Genotype Calls.” Nature Biotechnology 32 (3): 246–51. https://doi.org/10.1038/nbt.2835.

